# A conditional survival distribution-based method for censored data imputation: overcoming the hurdle in machine learning-based survival analysis

**DOI:** 10.1101/2022.02.18.481114

**Authors:** Yizhuo Wang, Christopher R. Flowers, Ziyi Li, Xuelin Huang

**Author notes:** **corresponding authors:** Xuelin Huang, 1400 Pressler Street, Unit 1411, Houston, TX 77030, Ziyi Li, 1400 Pressler Street, Unit 1410, Houston, TX 77030.

## Abstract

Data analyses by machine learning (ML) algorithms are gaining popularity in biomedical research. When time-to-event data are of interest, censoring is common and needs to be properly addressed. Most ML methods cannot conveniently and appropriately take the censoring information into consideration, potentially leading to inaccurate or biased results. We aim to develop a general-purpose method for imputing censored survival data, facilitating downstream ML analysis. In this study, we propose a novel method of imputing the survival times for censored observations. The proposal is based on their conditional survival distributions (CondiS) derived from Kaplan-Meier estimators. CondiS can replace censored observations with their best approximations from the statistical model, allowing for direct application of ML methods. When covariates are available, we extend CondiS by incorporating the covariate information through ML modeling (CondiS-X), which further improves the accuracy of the imputed survival time. Compared with existing methods with similar purposes, the proposed methods achieved smaller prediction errors and higher concordance with the underlying true survival times in extensive simulation studies. We also demonstrated the usage and advantages of the proposed methods through two real-world cancer datasets. The major advantage of CondiS is that it allows for the direct application of standard ML techniques for analysis once the censored survival times are imputed. We present a user-friendly R package to implement our method, which is a useful tool for ML-based biomedical research in this era of big data.

## 1. INTRODUCTION

Analyzing the time to the event of interest for patients with personalized predictors is a main goal of survival analysis. It provides valuable information about survival related factors and may enlighten directions for prolonging patients’ lives. Such analysis also assists clinicians in allocating resources more efficiently and developing suitable treatment plans for patients at different risk levels. More generally, survival analysis is a branch of statistics that involves the modelling of the time-to-event data.[1] Compared with traditional data analysis problems, partial observations of the outcome are more prevalent in survival analysis. This form of missing data issue is called censoring.[2] There are three types of censoring: left, right, and interval. Right-censoring, the most common, usually happens when patients drop out from the study or when the study ends earlier than the occurrence of the event of interest. Here, we focus on addressing right censoring in survival data, but the principal may also apply for other types of censoring events.

Given the importance of survival data, there is a long history of statistical method development to appropriately address censoring in survival analysis. These statistical approaches can be roughly divided into three categories: nonparametric methods (e.g., the Kaplan-Meier estimator[3] and the Nelson-Aalen estimator[4]), parametric methods (e.g., the Tobit model[5]), and semi-parametric methods (e.g., the Cox proportional hazards model[6]). The parametric and semi-parametric methods can be applied as regression-based models. However, most machine learning methods currently are developed for non-censored data. It takes huge amounts of efforts to re-develop current machine learning (ML) methods to handle censored data. In the literature, a lot of researchers chose to use ad-hoc techniques, which can lead to severely biased results. For example, some ignored the censoring information and directly applied ML methods for non-censored data to data with censoring, [7 8] while some simply discarded those censored observations.[9 10]

Currently, the healthcare industry has generated a vast amount of clinical data stored in electronic health records (EHRs) and genomic data created from laboratory experiments.[11] ML methods are gaining popularity as they are an advance in taking in this massive amount of data.[12] Several ML methods have been adapted to handle survival data, such as the survival tree model[13], the support vector approach[14], the neural network and the convolutional network methods [15] [16], as well as the deep survival and deep learning methods[17] [18]. However, all of these methods are reliant upon altering specific ML algorithms to handle censored data. The number of ML methods with the specific ability to handle censored data is limited. Thus, this adapting approach is not generalizable to the wide variety of existing ML methods. This limits the application of ML in biomedical research. Alternatively, researchers may simply discard censored observations or treat them as observed event times.[2] These ad hoc approaches ignore the nature of censoring events and may lead to biased or even wrong results. There is a need for statistically rigorous methods that can provide a general solution for addressing censored data in ML research.

The objective of this study is to develop a general method for right-censored time-to-event data for use in all ML algorithms. A popular imputation method proposed by Klein et al. generates pseudo-survival time based on right-censored data. [19] These pseudo values are derived from the imputation formula by comparing the whole population-averaged and leave-one-out estimators of the survival time.[20] Their method can be applied to all ML algorithms since the pseudo-observations are computed for all patients. However, there are a few limitations. First, the method imputes survival times even for those subjects whose survival times have been observed, and the imputed survival times usually are different from the truly observed survival times. Second, the Pseudo method heavily relies on a tuning parameter controlling the maximum follow up time. Inappropriate selection of this parameter may lead to poor results.

In this study, we proposed a novel method of imputing the survival times for censored observations based on their conditional survival distributions (CondiS) derived from the Kaplan-Meier estimator. CondiS can replace the censored observations with the best approximations from the statistical model, allowing for direct application of the ML methods. When covariates were available, we extended CondiS by incorporating the covariate information through ML modeling (CondiS-X), which further improved the survival time imputation. Using CondiS, only the observations of censored data were imputed.

Compared with the pseudo survival time method by Klein et al, our method is more natural. We studied the use of (1) raw data by treating censoring time as observed survival time (termed OBS), (2) the Pseudo method and (3) the CondiS method, on both simulated data and real-world data. When compared with the true survival time, using metrics including the concordance index, the mean absolute error (MAE), and the median absolute error (medAE), the simulation results show that CondiS outperforms the other two methods both before and after applying ML models. Similar conclusions were reached for real-data examples.

This paper is organized as follows: in Section 2, we first provide the statistical definition of the CondiS method and introduce its extension, CondiS-X. A schematic flowchart of our method is given. We also discuss the setup of the simulation design and some basic information of two real-world datasets. In addition, we describe three metrics to evaluate the performance of our method. In Section 3, we present the results of the Monte Carlo simulations and two empirical real-world examples. In Section 4, we illustrate the advantages and potential limitations of the proposed method. In Section 5, we briefly summarize this study. Additional proofs of the performance of our method are collected in the Supplementary Materials. We performed all the experiments in R and a user-friendly software package for implementing our proposed methods is available from https://github.com/yizhuo-wang/CondiS.

## 2. MATERIALS AND METHODS

The conventional survival models, such as the widely-used Cox model, usually are developed from the hazard function, which is defined as the instantaneous probability of the event of interest occurring within a narrow time frame.[1] For this reason, in traditional clinical trials the treatment effect frequently is reported in terms of the hazard ratio (HR).[21] HR is an estimator of relative risk/hazard rate reduction in different groups and can be obtained directly from a Cox proportional hazards model. However, many studies have shown that when the proportional hazards (PH) assumption of the treatment effect is violated, the estimation of the hazard rate is affected by the follow-up time in the study. This compromises the use of HR and makes it inaccurate.[22] Moreover, HR does not directly convert to differences in survival times, which makes it difficult to explain for non-statisticians. As a result, more interpretable measures are being explored for assessing the treatment effect.[22] Recently, the restricted mean survival time (RMST) has become popular for quantifying the between-group difference in survival analysis.[23] RMST does not depend on the proportional hazard assumption. It is a robust criterion for comparing treatments, and it is intuitively appealing to patients and physicians. Our proposed method, CondiS, is based on the concept of RMST and the time-conditioned survival function.

### CondiS

Suppose the Kaplan-Meier estimator of the survival function that the probability of life being longer than t is given by 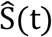. Then the mean survival time restricted to *t*^*^(RMST)[24] is:

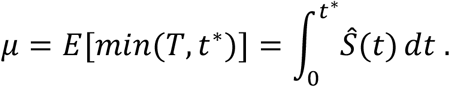

If patient *i* is censored at *t* = *t*_*i*_ < *t*^*^, then his expected survival time restricted to *t*^*^ can be estimated by a *t*^*\**^-year conditional survival distribution for patients who have survived for *t*_*i*_ years:

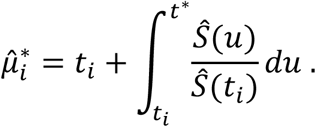

Suppose *t*_1_ ≤ *t*_2_ ≤ … ≤ *t*_*n*_, *t*^*^ can be any self-defined time-of-interest or can be defaulted as the largest survival time, *t*_*n*_. If *t*_*i*_ ⩾ *t*^*^, then 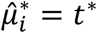 If patient *i* is not censored, then 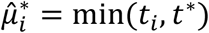 We propose to use 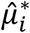 as an imputed survival time if patient *i’s* survival time is censored. After all censored survival times are imputed this way, they are combined with observed survival times and supplied to any ML method for analysis. Transformation may be applied to all the survival times (imputed or observed) before applying the ML analysis.

### CondiS-X

Ideally, we want the imputed survival time to be as close to the true survival time as possible. In the above imputation by CondiS, we did not take the covariate information into consideration. It is likely that some covariates are associated with patients’ survival and thus can help further improve our imputation. To use that covariate information, we extended CondiS by incorporating the covariates through ML modeling (CondiS-X). Specifically, we trained an ML model with the imputed or observed survival time as the dependent variable and covariates as independent variables. We then used this model to predict the censored survival times in the same dataset and used the predicted survival times to update the previously imputed values. This step may be done similar to cross-validation. That is, we randomly divide the whole dataset into *m* (e.g., *m*=10) partitions of equal sample size, use *m-1* partitions of the data to train the ML model, and use the resulting model to make survival time predictions for the remaining partition. We repeat this process for each of the *m* parts. A schematic diagram of the whole process of CondiS and CondiS-X is shown in Figure 1. A survival dataset including right-censored observations is our input data. We first apply CondiS to this dataset and obtain the imputed survival times for the censored observations. If the covariates information is available, we will add an extra step that uses CondiS-X to update the imputed survival times. Now we have a survival dataset without censoring that we can use to apply any ML technique.

**Figure 1.**
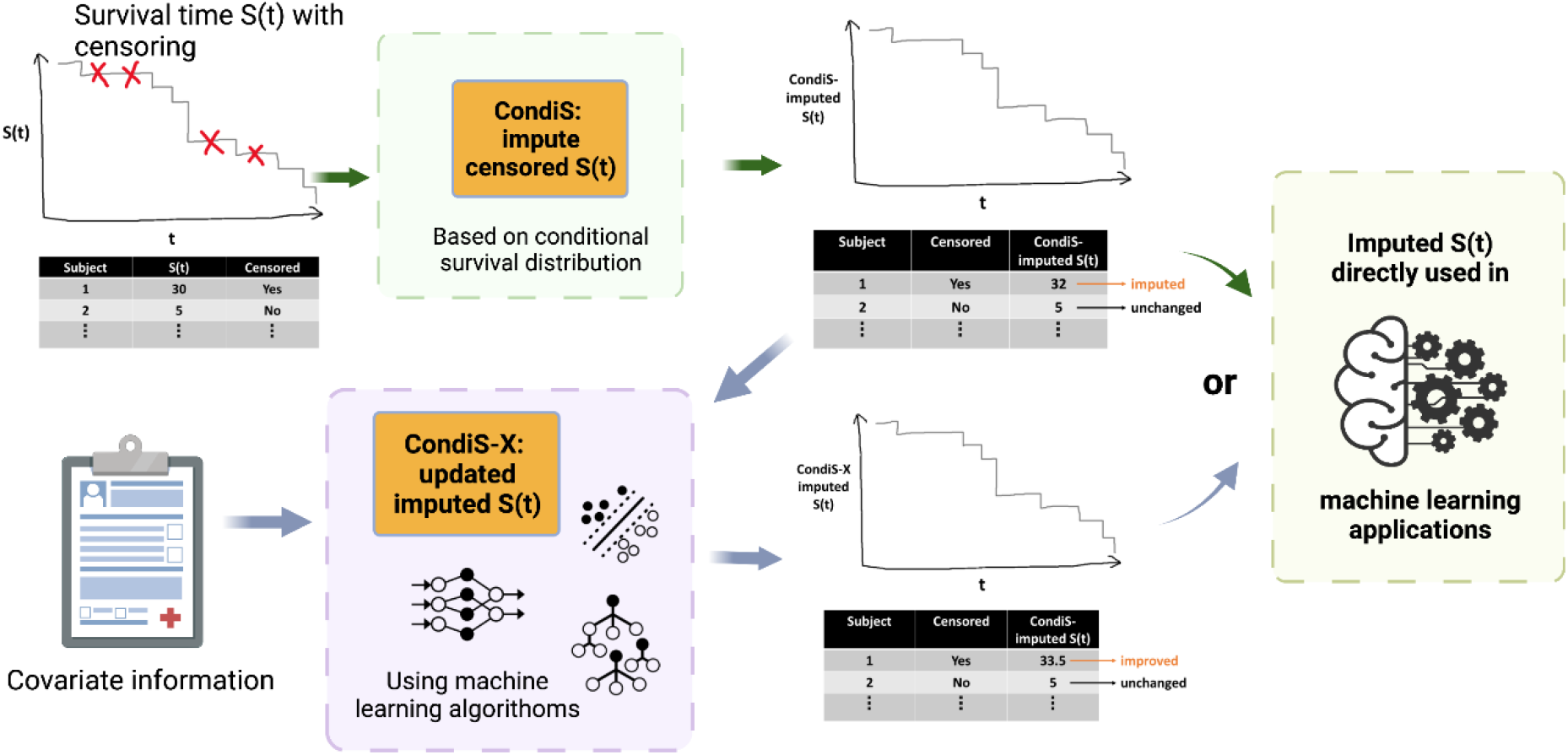
Schematic of the CondiS pipeline. Created with BioRender.com.

### Simulation design

We use the inverse probability method presented by Bender et al. to generate true survival times *T* from the proportional hazards model.[25] Let *S*(· | *x*) be the conditional survival function derived from the proportional hazards model:

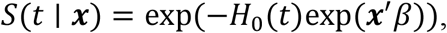

where *H*_0_(*t*) is the baseline cumulative hazard function, *x* is the given covariate vector and *β* is the regression coefficient. Generate random numbers *W* from a uniform distribution on [0,1], then we can generate *T* through

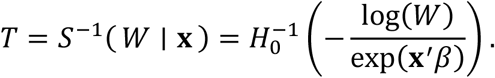

Here we assume that at the baseline, *T* follows a Weibull distribution with a shape parameter *ρ* > 0 and a scale parameter *λ* > 0, i.e., we have the cumulative hazard function and its inverse:

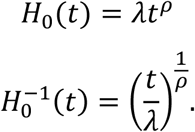

Then we generate the Weibull survival time conditional on covariates *x* by drawing

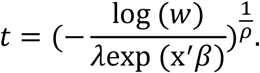

In this example, the covariate matrix consists of ten biomarkers, one binary treatment, two treatment-biomarker interaction terms and one random noise:

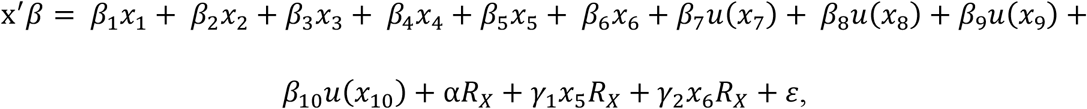

where *x*_1_∼*x*_10_ are biomarkers generated from a normal distribution with a mean of 0 and a standard deviation of 1, *R*_*X*_ is a binary treatment with either 0 or 1, *β*_1_∼*β*_10_ are biomarkers’ coefficients, α is the treatment main effect coefficient, *γ*_1_∼*γ*_2_ are the treatment-biomarker interaction coefficients, and *ε* is a tunable random noise. Among 10 biomarkers, *x*_7_∼*x*_10_ contribute to the Weibull survival time as step functions:

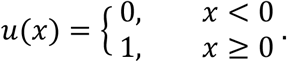

To create right-censored observations, we drew random times from another Weibull distribution as the censoring time *C*:

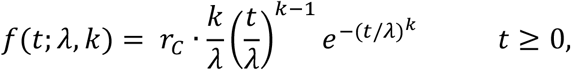

where *k* > 0 is the shape parameter, *λ* > 0 is the scale parameter of the distribution, and *r*_*C*_ is a random number to tune the censoring time *C*. Then our final observed survival time *Y* can be computed as *Y* = *min*(*T, C*).

We applied the Pseudo and CondiS methods and obtained their survival times. The better method should be closer to the true survival time. For the extension, CondiS-X, we built 7 commonly used ML models, including the generalized linear model (GLM)[26], LASSO regression[27], Ridge regression[28], gradient boosting machine (GBM)[29], random forest (RF)[30], support vector machine (SVM)[31], k-nearest neighbors (KNN)[32], and artificial neural networks (ANN)[33]. Note that the same models were built using the raw observed time (OBS-X) and the pseudo time (Pseudo-X) for later comparison.

### Real-world examples

We applied our method to two real-world datasets. The first one included 2982 primary breast cancers patients whose data were included in the Rotterdam tumor bank.[34] We took only the uncensored part of the data, which left us with 1214 observations. We used only the observed survival time so that we had a gold standard to evaluate our methods. For the dataset of 1214 observations, we artificially generated censoring times *C* from a uniform distribution. The final observed survival time *Y* was computed as *Y* = *min*(*T, C*), where *T* was the true survival time from the original Rotterdam dataset. In this way, we were able to evaluate the performance of our method after applying it to the dataset of 1214 observations, some of which were censored by the artificially generated censoring times *C*. The second dataset we used was a blood cancer dataset, which included a cohort of 1,001 diffuse large B cell lymphoma (DLBCL) patients. [35] We kept the 574 uncensored subjects and, similarly, we simulated censoring times for this DLBCL dataset with 574 observations to apply and test our method.

### Evaluation metrics

For censored data, standard evaluation metrics for regression problems such as the root of the mean squared error and R^2^ are not appropriate for measuring the performance in survival analysis.[36] In our study, we resorted to other evaluation metrics. One of the most frequently used metrics in survival analysis is the concordance index (CI), which is a measure of the rank correlation.[37] We used CI to assess the correlation between imputed survival time and the true survival time. Given the comparable instance pair (*i, j*) with *T*_*i*_ and *T*_*j*_ are the true survival times and 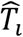 and 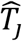 are the imputed survival times, the concordance index *C* measures the probability of concordance between the rankings of actual values and imputed values:

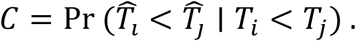

Mean absolute error (MAE) is another commonly-used evaluation metric, which measures the average magnitude of errors between paired observations. [38] The formula is

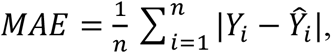

where *Y*_*i*_ is the true survival time and *Ŷ*_*i*_ is the simulated survival time.

Median absolute error (medAE) is also used here because it is robust to outliers.[38] It is calculated by taking the median of all absolute differences between the true survival time and the simulated survival time.

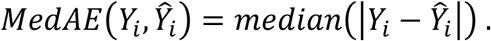

Using these three metrics, we thoroughly compared the CondiS imputed survival time to the Pseudo survival time. Meanwhile, since we extended our method by incorporating covariate information through ML models (CondiS-X), we should also do the same thing to other methods when comparing them. We extended the raw observation time (OBS), i.e., ignoring the censoring status, and the Pseudo survival time to OBS-X and Pseudo-X through the same ML models. Then we compared the performance of OBS-X, Pseudo-X, and CondiS-X using CI, MAE, and medAE.

## 3. RESULTS

### Simulation

Five scenarios of different censoring rates (% censoring = 20, 30, 40, 50, 60) were considered. We conducted 1000 Monte Carlo simulations for each scenario and compared the results of the raw observed time, the Pseudo method, and the CondiS method. Since MAE may be greatly distorted by extreme outliers, we also present medAE. CI describes the consistency between observed and predicted relative ranks of the survival time of the subjects. For MAE and medAE, smaller is better. For CI, larger is better.

In Figure 2, (A), (B), and (C) show the results of MAE, medAE, and CI comparing the CondiS and the Pseudo method. The x-axis is the censoring percentage; the Pseudo method and CondiS method are represented by the grey box and yellow box, respectively. A lower MAE value in

**Figure 2.**
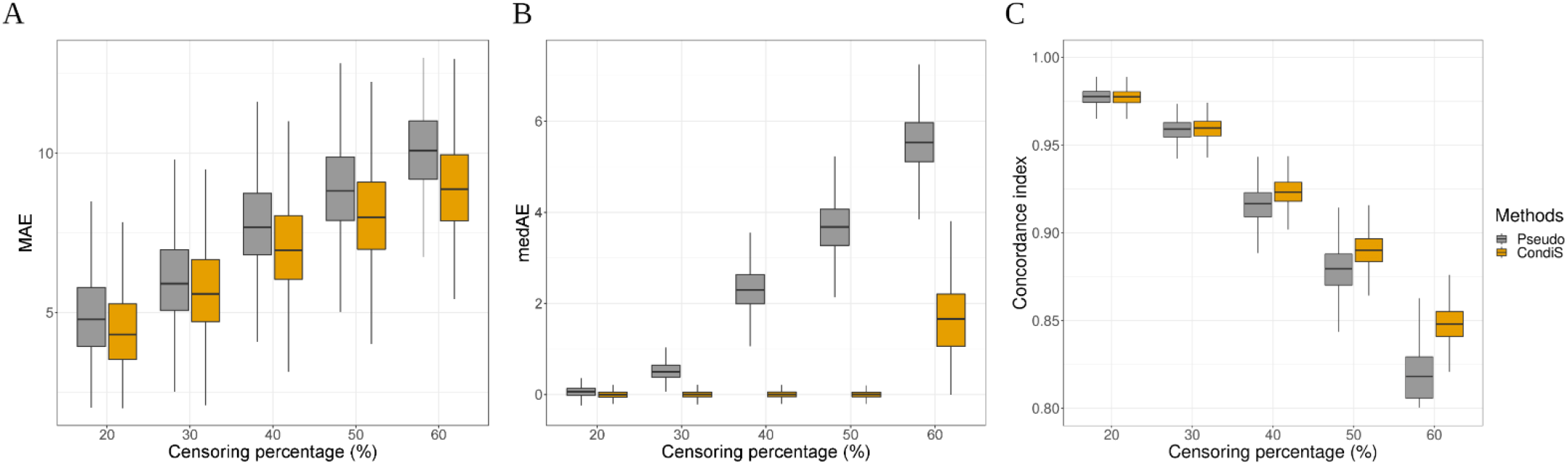
Simulation study: evaluation of the Pseudo method and the CondiS method at different censoring percentages. (A) Mean absolute error of the imputed survival time and true survival time. (B) Median absolute error of the imputed survival time and true survival time. (C) Concordance index of the imputed survival time and true survival time. Boxplots display the median (middle line), the inter-quartile range (hinges), and 1.5 times the inter-quartile range (lower and upper whiskers) based on a 1000-time simulation.

Figure 2(A) and a higher CI value in Figure 2(C) indicate that the imputed survival time by the CondiS method is more similar to the true survival time than to that of the Pseudo method. The advantage of CondiS over Pseudo holds across different censoring rate levels. For higher censoring rate levels, the difference between CondiS imputed survival time and the Pseudo-survival time becomes even larger. In Figure 2(B), we can see that the medAE value remains at zero until the censoring percentage raises to 60%. This makes sense because, using the CondiS method, only the censored observations in the dataset are replaced. When the censoring percentage is less 50%, the medAE is always zero because more than 50% of survival times are observed, and they stay the same before and after applying the CondiS method, resulting in zero absolute errors.

### CondiS-X

To improve the imputed survival time, we utilized the given covariate information through different ML models. Fixing the censoring rate at 40%, the results of MAE, medAE, and CI comparing OBS-X, Pseudo-X, and CondiS-X are shown in Figure 3 (A), (B), and (C). Again, in Figure 3(A) and Figure 3(C), CondiS X gives us the lowest MAE and the highest CI among these three methods. And as we would expect, the medAE values of CondiS-X remain at zero for each of the ML models because the censoring percentage is under 50%. Furthermore, we can see that, when X is a nonparametric model such as the GBM, the RF, and the SVM models, we obtain better results. For this specific simulation dataset, these models maximize the effect of covariates and improve the imputed time to a greater extent.

**Figure 3.**
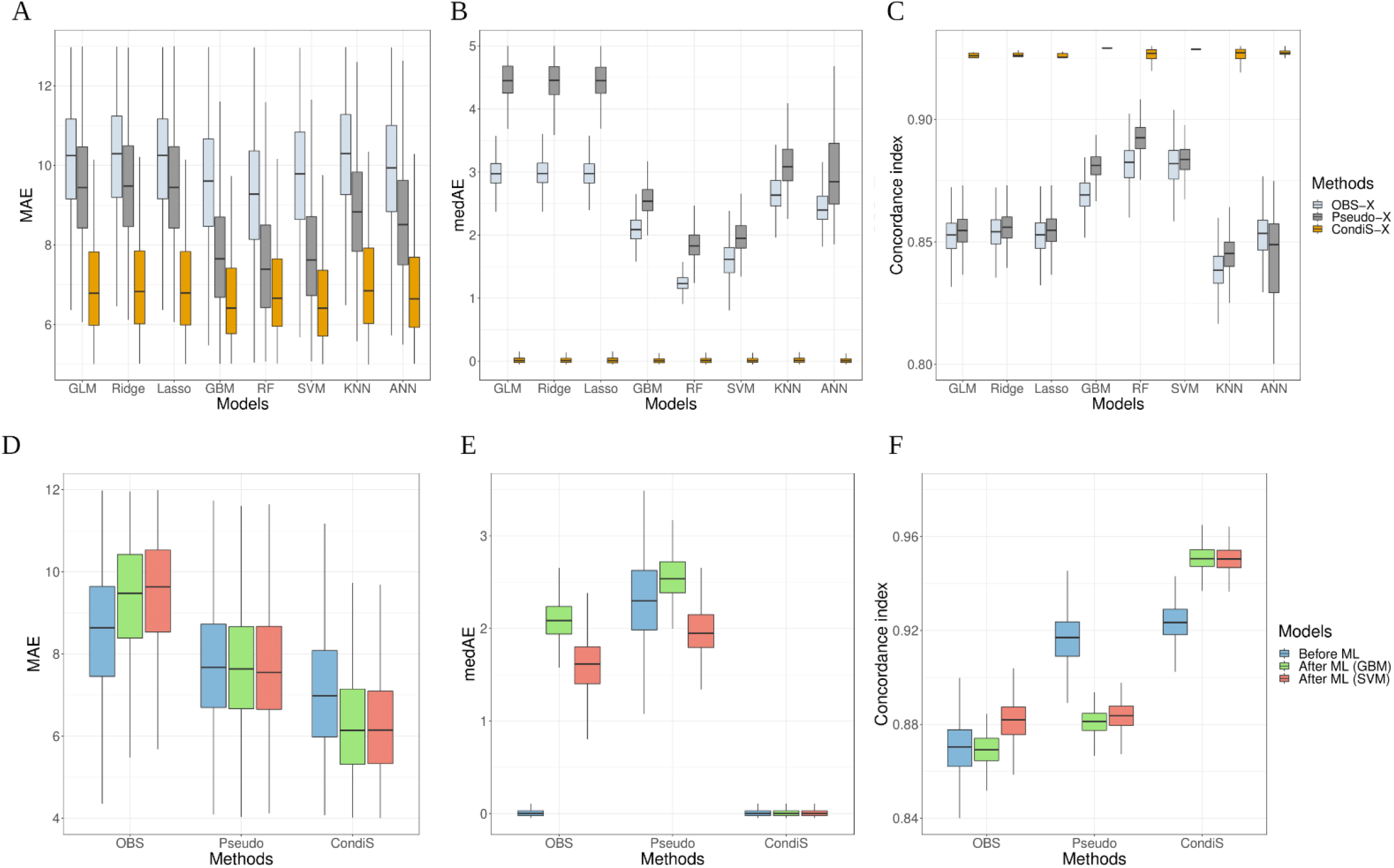
Simulation study: evaluation of the OBS-X method, the Pseudo-X method, and the CondiS-X method at 40% censoring. (A, B, C) Mean absolute error, median absolute error, and concordance index of ML-improved survival time and true survival time, respectively. (D, E, F) Comparison of before ML-extension performance and after ML-extension performance for OBS, Pseudo, and CondiS methods, respectively. Boxplots display the median (middle line), the inter-quartile range (hinges), and 1.5 times the inter-quartile range (lower and upper whiskers) based on a 1000-time simulation.

In Figure 3(D), (E), and (F) we compared the performance of each method before and after the ML extension by the GBM and SVM algorithms. We also evaluated which method benefited the most from this extra step. Regarding the MAE in Figure 3(D), the performance of the OBS method is worse, the performance of the Pseudo method does not show an obvious boost, and only the CondiS method resulted in a lower error. Regarding the medAE in Figure 3(E), the CondiS method maintained its advantage of a zero median error. Regarding the CI in Figure 3(F), the performance of the OBS method was unstable, the performance of the Pseudo method was worse, and only the CondiS method improved correlation with the true survival time. In summary, we conclude that the CondiS method benefited from the covariate information and improved its imputed survival time.

### Real-world examples

We tested CondiS/CondiS-X on two real-world datasets, the Rotterdam dataset [34] and the DLBCL dataset.[35] Similarly, we compared the performance of our method to the OBS and the Pseudo methods. We took only the uncensored subjects with true survival times and then simulated censoring times for these subjects. In this way, we have a gold standard to evaluate our methods.

For both real-world datasets, we observed a similar pattern as for the simulation. Figure 4 (A), (B), and (C) show the results of the Rotterdam dataset, and Figure 6 (A), (B), and (C) show the results of the DLBCL dataset. In these figures, CondiS is represented by the yellow line and Pseudo is represented by the grey line. We can see that, for each different censoring percentage, compared with Pseudo, the imputed time of CondiS has low MAE, medAE values and a high CI value. In other words, it is much closer to the true survival time and has a higher correlation with the true survival time, too.

**Figure 4.**
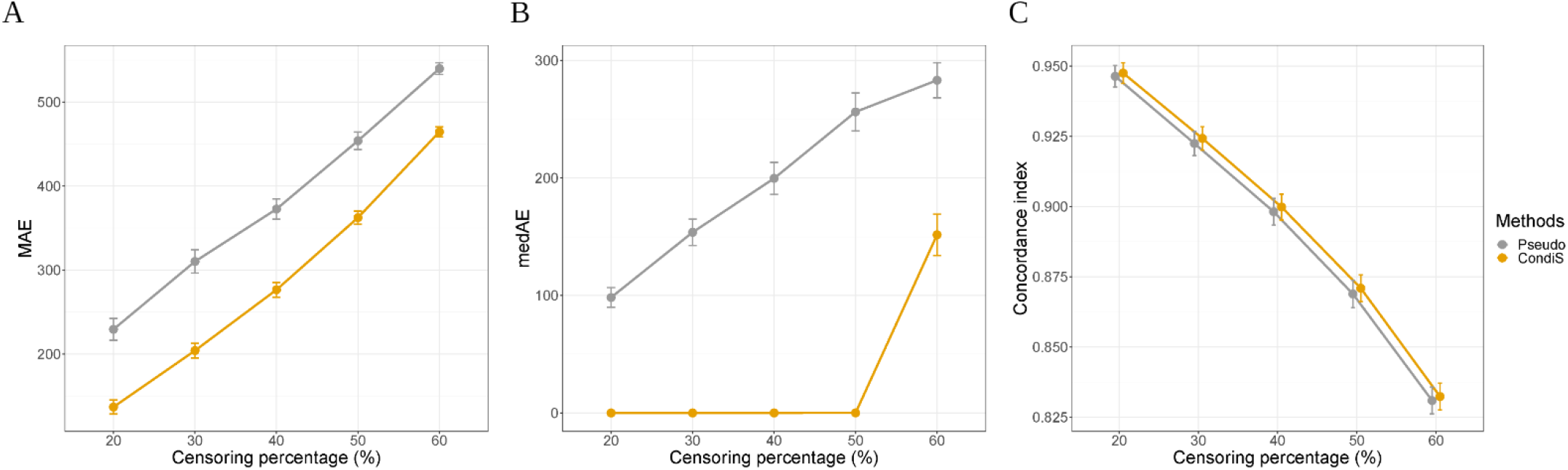
Rotterdam dataset: evaluation of the Pseudo method and the CondiS method at different censoring percentages. (A) Mean absolute error of the imputed survival time and true survival time. (B) Median absolute error of the imputed survival time and true survival time. (C) Concordance index of the imputed survival time and true survival time. Line plots display the confidence interval based on 500 repetitions of the simulated censoring time.

For the Rotterdam dataset, we have done the extra ML modeling step to improve the imputed survival time and the results are shown in Figure 5. From Figure 5 (A), (B), and (C), we clearly observe a definite advantage of the CondiS method for every ML model. For this specific dataset, in terms of narrowing the gap between the imputed survival time and the true survival time, parametric algorithms on the left side of the figures perform better overall than non-parametric methods on the right side of the figures. We selected the Ridge and the Lasso algorithms to compare before and after extension performance for all three methods (OBS-X, Pseudo-X, CondiS-X) and the results are shown in Figure 5 (D), (E), and (F). The OBS-X method is getting further away from the true survival times; the Pseudo-X method achieves improved results; however, the CondiS method displays greater improvement and has the best performance.

**Figure 5.**
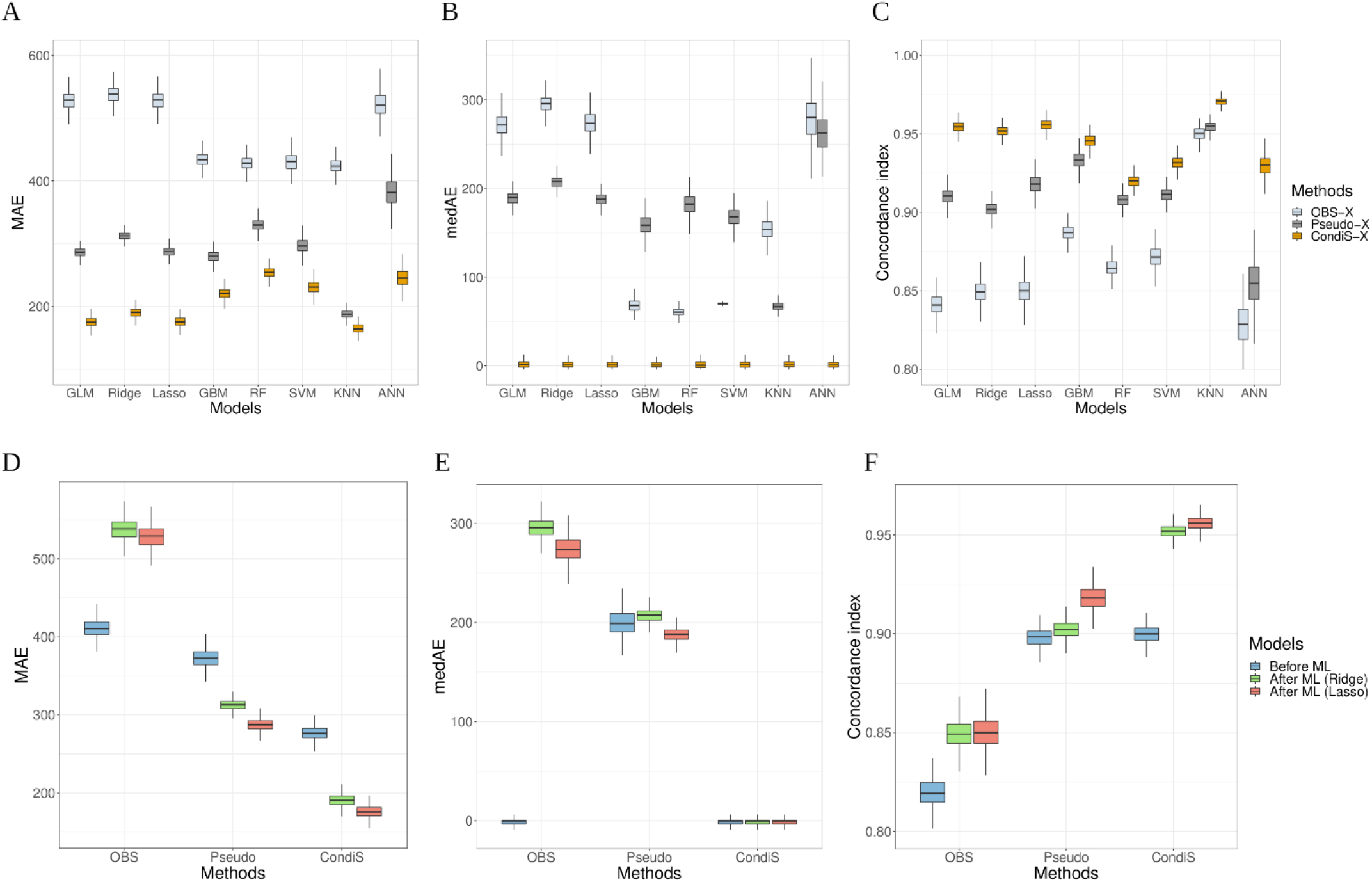
Rotterdam dataset: evaluation of the OBS-X method, the Pseudo-X method, and the CondiS-X method at 40% censoring. (A, B, C) Mean absolute error, median absolute error and concordance index of ML-improved survival time, and true survival time, respectively. (D, E, F) Comparison of before ML performance and after ML performance for OBS, Pseudo, and CondiS methods, respectively. Boxplots display the median (middle line), the inter-quartile range (hinges), and 1.5 times the inter-quartile range (lower and upper whiskers) based on 500 repetitions of the simulated censoring time.

**Figure 6.**
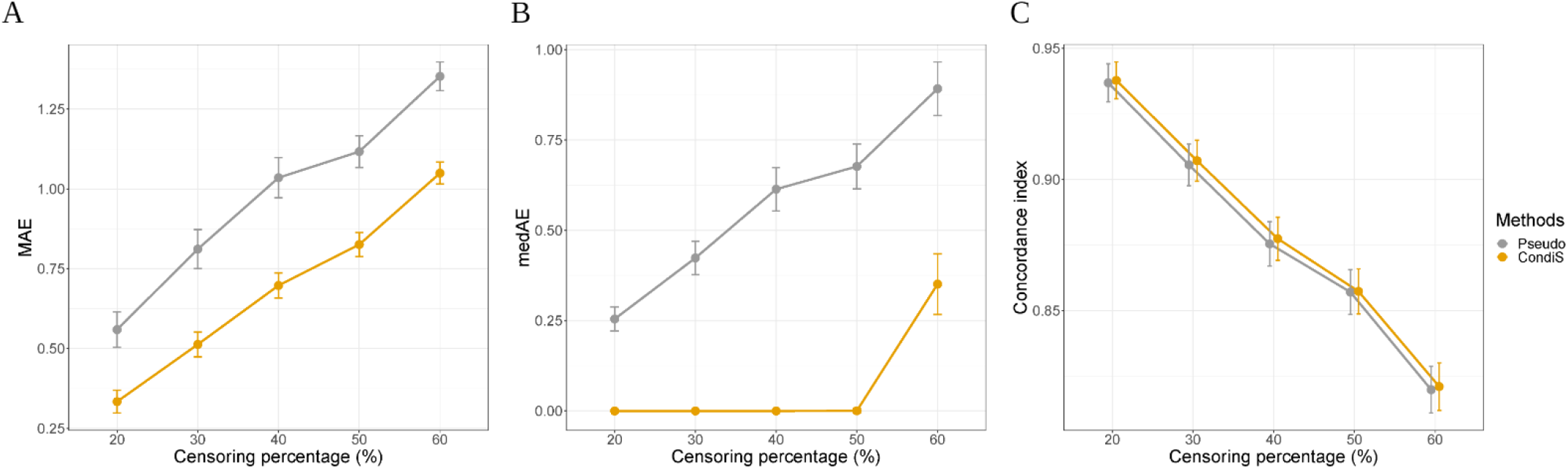
DLBCL dataset: evaluation of the Pseudo method and the CondiS method at different censoring percentages. (A) Mean absolute error of the imputed survival time and true survival time. (B) Median absolute error of the imputed survival time and true survival time. (C) Concordance index of the imputed survival time and true survival time. Line plots display the confidence interval based on 500 repetitions of the simulated censoring time.

## 4. Discussion

This study introduced a method (CondiS) to impute censored survival data, allowing for direct application to all ML models. Moreover, we extended our method by utilizing the covariate information through ML models. Regarding which ML algorithm to use, it depends on the data. For example, if there are lots of non-linear relationships and interactions in the data, one of the nonparametric models might be the appropriate choice. No matter which algorithm is used, the imputed time will always get closer to the true survival time as long as your covariates are relatively significant in the model.

The biggest advantage of our proposed method is its convenience. It is a one-method-fits-all approach applicable to all ML methods. It is not intended for optimal efficiency of the estimations. Rather, for each ML method there are likely methods tailored for that specific method that can achieve better performance. Our future research includes inspecting each ML method to propose more efficient methods for handling time-to-event data with censoring.

Both the simulation and the real-world examples’ results validate the advantages of our method over other methods with similar purposes. At different censoring percentage ranges from 20% to 60%, the imputed time of the CondiS methods is always the most similar to the true survival time in terms of low MAE, medAE values, and a high CI value. Aside from changing the censoring rate, we also evaluate our method by changing a self-defined parameter (t*) in our statistical model. For the Klein et al. Pseudo method, they also have a self-defined parameter, tmax, which is the maximum cut-off point for the restricted mean survival time. This parameter is comparable to the t* in our method. Both parameters help to prespecify a follow-up period that is of interest. We took a step further to demonstrate the advantage of our method by comparing these two methods under the same tmax/t*. The MAE, medAE, and CI results of the simulation and two real-world examples are plotted in Supplementary Figures S1-S3, respectively. We can see that, at a fixed censoring rate of 40% for the same prespecified time parameter, the imputed time of the CondiS method has more similarities with the true survival time than that of the Pseudo method. Given that many ML approaches drop individuals where censoring occurs, applying the CondiS method to the Rotterdam and DLBCL datasets would increase the effective sample size for these real-world datasets.

## 5. Conclusion

Censoring complicates the simple regression analysis of survival time and also causes inconvenience when applying complex ML algorithms. In this study, we have proposed an alternative method to impute the censored data based on the conditional survival distribution (CondiS). Compared with existing methods with similar objectives, the imputed time of our method is able to be more like the underlying true survival time by achieving a smaller mean absolute error, a smaller median absolute error, and a higher concordance index. When covariates are available, we further improve our method by utilizing the covariate information through different machine learning models (CondiS-X), which helps to predict the survival time more accurately. The major advantage of our method is that it allows for direct use of standard ML techniques for analysis once the imputed survival time is obtained.

## SUPPLEMENTARY FILES

**Figure S1:**
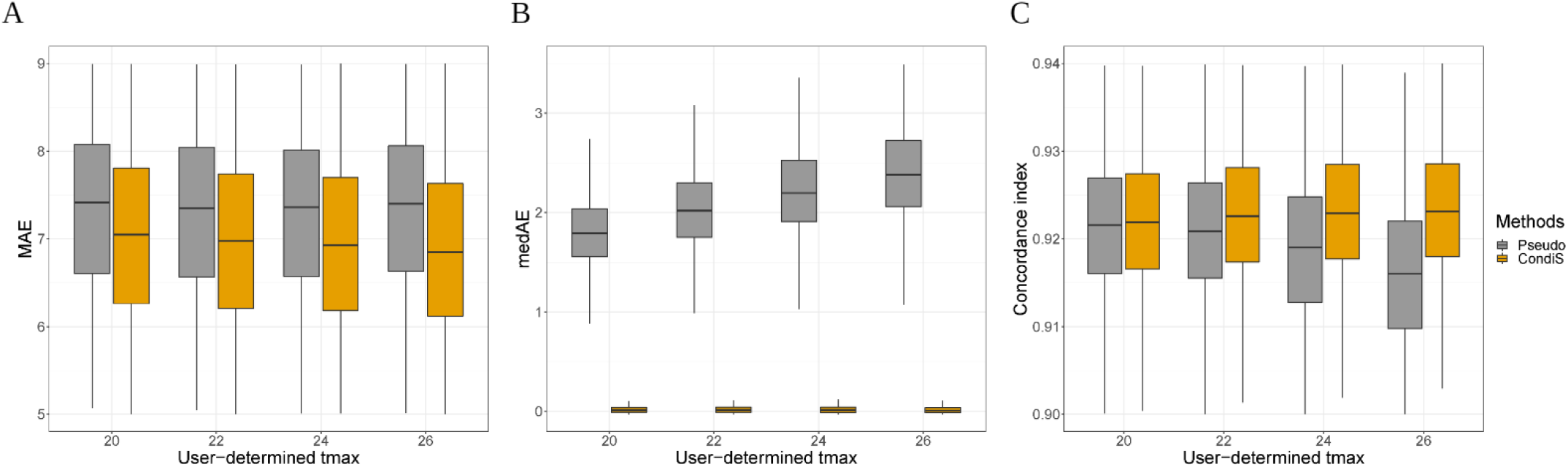
Simulation study: evaluation of the Pseudo method and the CondiS method with the same user-defined time parameter at 40% censoring. (A) Mean absolute error of the imputed survival time and true survival time. (B) Median absolute error of the imputed survival time and true survival time. (C) Concordance index of the imputed survival time and true survival time. Boxplots display the median (middle line), the inter-quartile range (hinges), and 1.5 times the inter-quartile range (lower and upper whiskers) based on a 1000-time simulation.

**Figure S2:**
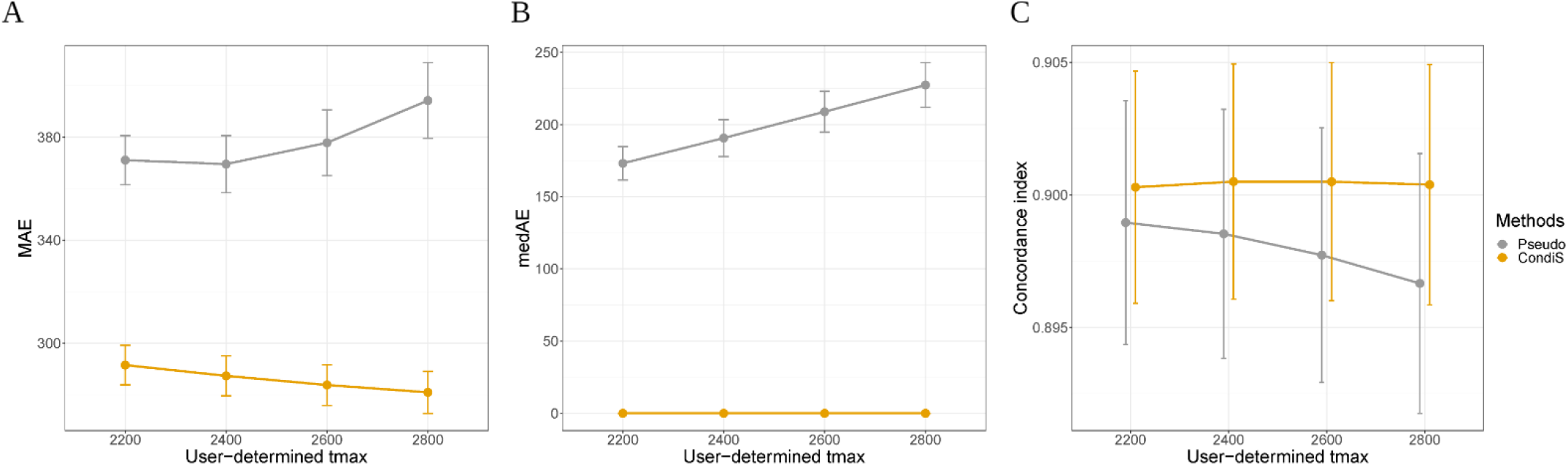
Rotterdam dataset: evaluation of the Pseudo method and the CondiS method with the same user-defined time parameter at 40% censoring. (A) Mean absolute error of the imputed survival time and true survival time. (B) Median absolute error of the imputed survival time and true survival time. (C) Concordance index of the imputed survival time and true survival time. Line plots display the confidence interval based on 500 repetitions of the simulated censoring time.

**Figure S3:**
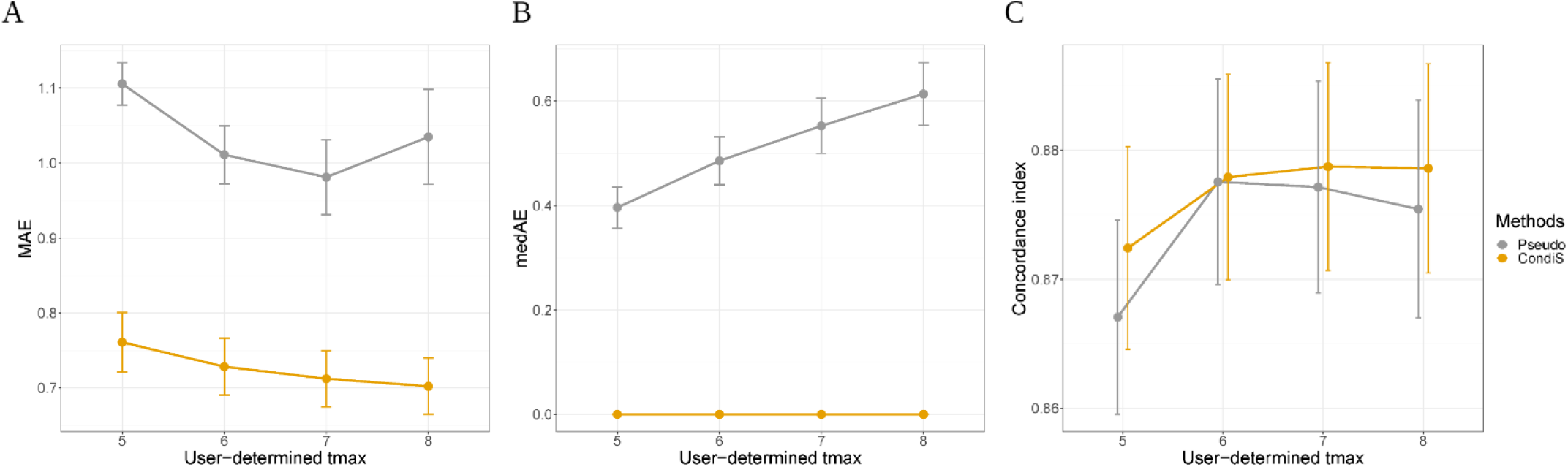
DLBCL dataset: evaluation of the Pseudo method and the CondiS method with the same user-defined time parameter at 40% censoring. (A) Mean absolute error of the imputed survival time and true survival time. (B) Median absolute error of the imputed survival time and true survival time. (C) Concordance index of the imputed survival time and true survival time. Line plots display the confidence interval based on 500 repetitions of the self-simulating censoring time.

## AUTHOR CONTRIBUTIONS

XH, ZL and YW conceived the concept of the study and designed the method. YW implemented the method, performed the simulations, and drafted the initial manuscript. CF provided guidance on interpreting the real-world data. All authors edited and approved the final manuscript.

## DATA AVAILABILITY STATEMENT

The Rotterdam dataset is a built-in R dataset called *rotterdam* and the DLBCL dataset can be downloaded from: https://www.ncbi.nlm.nih.gov/pmc/articles/PMC5659841/#SD2.

## CODE AVAILABILITY STATEMENT

An R package to implement the proposed method can be installed from https://github.com/yizhuo-wang/CondiS.

---

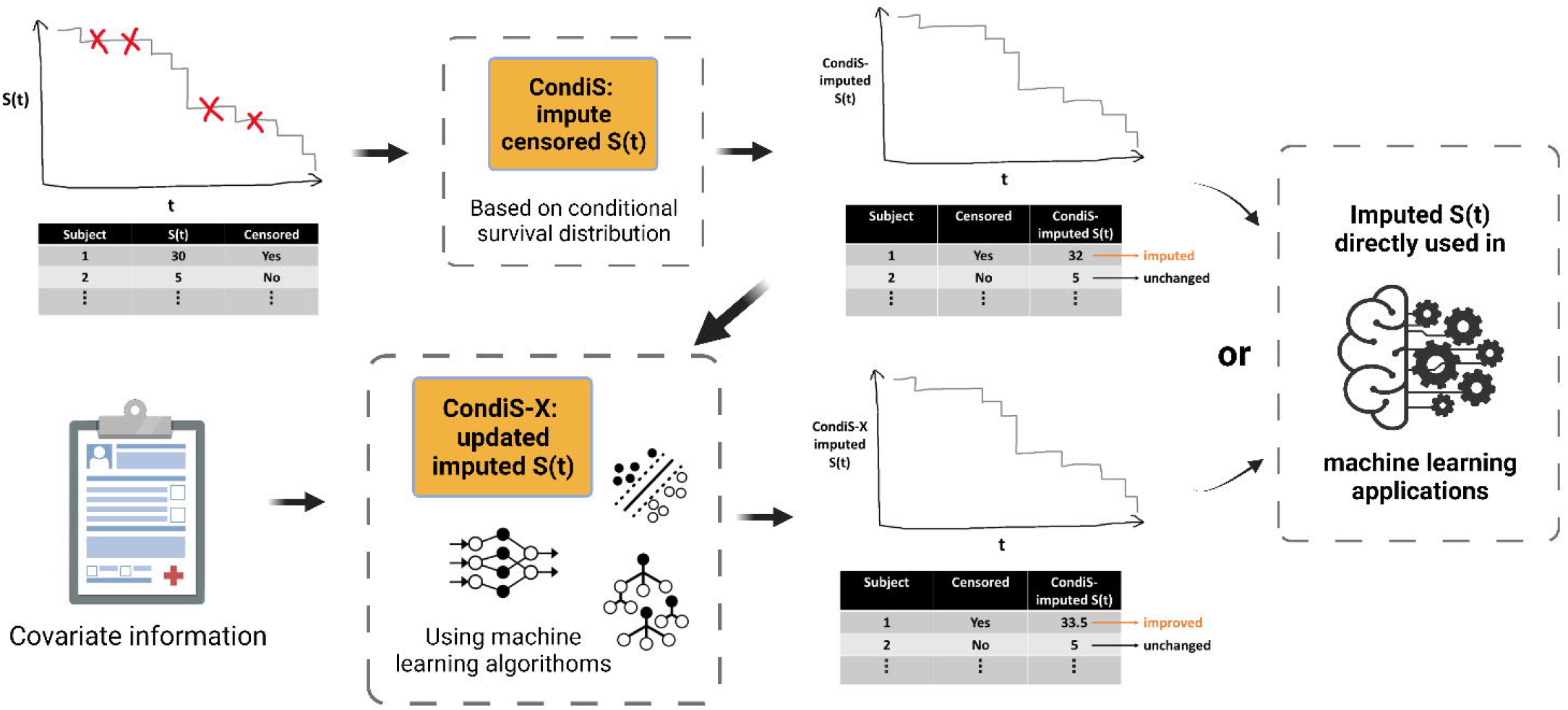

## REFERENCES

1. Singh R, Mukhopadhyay K. Survival analysis in clinical trials: Basics and must know areas. Perspect Clin Res 2011;2(4):145–8.

2. Prinja S, Gupta N, Verma R. Censoring in clinical trials: review of survival analysis techniques. Indian J Community Med 2010;35(2):217–21.

3. Kaplan EL, Meier P. Nonparametric Estimation from Incomplete Observations. Journal of the American Statistical Association 1958;53(282):457–81.

4. Nelson W. Hazard Plotting for Incomplete Failure Data. Journal of Quality Technology 1969;1(1):27–52.

5. Tobin J. Estimation of Relationships for Limited Dependent Variables. Econometrica 1958;26(1):24–36.

6. Cox DR. Regression Models and Life-Tables. Journal of the Royal Statistical Society. Series B (Methodological) 1972;34(2):187–220.

7. Kattan MW, Hess KR, Beck JR. Experiments to determine whether recursive partitioning (CART) or an artificial neural network overcomes theoretical limitations of Cox proportional hazards regression. Computers and biomedical research 1998;31(5):363–73.

8. Štajduhar I, Dalbelo-Bašić B, Bogunović N. Impact of censoring on learning Bayesian networks in survival modelling. Artificial intelligence in medicine 2009;47(3):199–217.

9. Sierra B, Larranaga P. Predicting survival in malignant skin melanoma using Bayesian networks automatically induced by genetic algorithms. An empirical comparison between different approaches. Artificial intelligence in Medicine 1998;14(1-2):215–30.

10. Blanco R, Inza I, Merino M, Quiroga J, Larrañaga P. Feature selection in Bayesian classifiers for the prognosis of survival of cirrhotic patients treated with TIPS. Journal of Biomedical Informatics 2005;38(5):376–88.

11. Raghupathi W, Raghupathi V. Big data analytics in healthcare: promise and potential. Health Inf Sci Syst 2014;2:3.

12. Dasgupta A, Sun YV, Konig IR, Bailey-Wilson JE, Malley JD. Brief review of regression-based and machine learning methods in genetic epidemiology: the Genetic Analysis Workshop 17 experience. Genet Epidemiol 2011;35 Suppl 1:S5–11.

13. Leblanc M, Crowley J. Survival Trees by Goodness of Split. Journal of the American Statistical Association 1993;88(422):457–67.

14. A Support Vector Approach to Censored Targets. Seventh IEEE International Conference on Data Mining (ICDM 2007); 2007 28-31 Oct. 2007.

15. Faraggi D, Simon R. A neural network model for survival data. Statistics in Medicine 1995;14(1):73–82.

16. Mobadersany P, Yousefi S, Amgad M, et al. Predicting cancer outcomes from histology and genomics using convolutional networks. Proceedings of the National Academy of Sciences 2018;115(13):E2970–E79.

17. Katzman J, Shaham U, Cloninger A, Bates J, Jiang T, Kluger Y. Deep Survival: A Deep Cox Proportional Hazards Network. ArXiv 2016;abs/1606.00931.

18. Yousefi S, Amrollahi F, Amgad M, et al. Predicting clinical outcomes from large scale cancer genomic profiles with deep survival models. Scientific Reports 2017;7(1):11707.

19. Klein JP, Gerster M, Andersen PK, Tarima S, Perme MP. SAS and R functions to compute pseudo-values for censored data regression. Comput Methods Programs Biomed 2008;89(3):289–300.

20. Andersen PK, Hansen MG, Klein JP. Regression Analysis of Restricted Mean Survival Time Based on Pseudo-Observations. Lifetime Data Analysis 2004;10(4):335–50.

21. Spruance SL, Reid JE, Grace M, Samore M. Hazard ratio in clinical trials. Antimicrob Agents Chemother 2004;48(8):2787–92.

22. Keene ON. Alternatives to the hazard ratio in summarizing efficacy in time-to-event studies: an example from influenza trials. Stat Med 2002;21(23):3687–700.

23. Royston P, Parmar MK. The use of restricted mean survival time to estimate the treatment effect in randomized clinical trials when the proportional hazards assumption is in doubt. Stat Med 2011;30(19):2409–21.

24. Royston P, Parmar MK. Restricted mean survival time: an alternative to the hazard ratio for the design and analysis of randomized trials with a time-to-event outcome. BMC Med Res Methodol 2013;13:152.

25. Bender R, Augustin T, Blettner M. Generating survival times to simulate Cox proportional hazards models. Stat Med 2005;24(11):1713–23.

26. Nelder JA, Wedderburn RWM. Generalized Linear Models. Journal of the Royal Statistical Society. Series A (General) 1972;135(3):370–84.

27. Tibshirani R. Regression Shrinkage and Selection via the Lasso. Journal of the Royal Statistical Society. Series B (Methodological) 1996;58(1):267–88.

28. Hoerl AE, Kennard RW. Ridge Regression: Biased Estimation for Nonorthogonal Problems. Technometrics 1970;12(1):55–67.

29. Friedman JH. Greedy function approximation: A gradient boosting machine. The Annals of Statistics 2001;29(5):1189–232, 44.

30. Ho TK. Random decision forests. Proceedings of the Third International Conference on Document Analysis and Recognition (Volume 1) - Volume 1: IEEE Computer Society, 1995:278.

31. Aizerman MA. Theoretical Foundations of the Potential Function Method in Pattern Recognition Learning. Automation and Remote Control 1964;25:821–37.

32. Altman NS. An Introduction to Kernel and Nearest-Neighbor Nonparametric Regression. The American Statistician 1992;46(3):175–85.

33. Grossberg S. Nonlinear neural networks: Principles, mechanisms, and architectures. Neural Networks 1988;1(1):17–61.

34. Royston P, Altman DG. External validation of a Cox prognostic model: principles and methods. BMC Med Res Methodol 2013;13:33.

35. Reddy A, Zhang J, Davis NS, et al. Genetic and Functional Drivers of Diffuse Large B Cell Lymphoma. Cell 2017;171(2):481–94 e15.

36. Heagerty PJ, Zheng Y. Survival Model Predictive Accuracy and ROC Curves. Biometrics 2005;61(1):92–105.

37. Uno H, Cai T, Pencina MJ, D’Agostino RB, Wei LJ. On the C-statistics for evaluating overall adequacy of risk prediction procedures with censored survival data. Stat Med 2011;30(10):1105–17.

38. Hyndman R. Another Look at Forecast Accuracy Metrics for Intermittent Demand. Foresight: The International Journal of Applied Forecasting 2006;4:43–46.

